# Converging evidence that left extrastriate body area supports visual sensitivity to social interactions

**DOI:** 10.1101/2023.05.23.541943

**Authors:** Marco Gandolfo, Etienne Abassi, Eva Balgova, Paul E. Downing, Liuba Papeo, Kami Koldewyn

## Abstract

Navigating our complex social world requires processing the interactions we observe. Recent psychophysical and neuroimaging studies provide parallel evidence that the human visual system may be attuned to efficiently perceive dyadic interactions. This work implies, but has not yet demonstrated, that activity in body-selective cortical regions causally supports efficient visual perception of interactions. We adopt a multi-method approach to close this important gap. First, using a large fMRI dataset (N=92), we found that the left-hemisphere Extrastriate Body Area (EBA) responds more to face-to-face than non-facing dyads. Second, we replicated a behavioural marker of visual sensitivity to interactions: categorisation of facing dyads is more impaired by inversion than non-facing dyads. Third, in a pre-registered experiment, we used fMRI-guided transcranial magnetic stimulation to show that online stimulation of the left EBA, but not a nearby control region, abolishes this selective inversion effect. Activity in left EBA, thus, causally supports the efficient perception of social interactions.

## Introduction

As a social species, humans are constantly engaged in interactions with each other. This places strong demands on neural systems that must make sense of others’ behaviour and attempt to infer underlying emotional and mental states as well as discern longstanding traits (Adolphs, 2001, 2009; Brothers, 1990). While we stand to learn much from our own direct interactions with others, many valuable social cues are also found in observing the interactions between other individuals. Such observed interactions can carry unique cues about others’ relationships, social skills, goals, and personalities. Accordingly, in contrast to much of the “social vision” work on perception of social cues from individual faces, bodies, and voices (e.g., Adams et al., 2011; Belin et al., 2011; Duchaine & Yovel, 2015; Hu et al., 2020; Peelen & Downing, 2007), researchers have recently turned their attention to the visual processes underpinning perception of interactions per se.

One recent perspective is that facing pairs (dyads) of humans elicit an emergent, holistic percept, one that is more than the sum of its component parts, and that may engage unique perceptual processes. This proposal is supported, for example, by findings that facing dyads are particularly prone to the disrupting effect of in-plane inversion (Papeo, 2020; Papeo et al., 2017), similar to other socially-relevant stimuli such as individual faces (Yin, 1969) and bodies (Gandolfo & Downing, 2020; Griffin & Oswald, 2022; Reed et al., 2003). Indeed, stimulus inversion strongly impairs recognition performance of social stimuli, compared to other objects. When a social stimulus (e.g. a face) is presented in its typical, upright position it is readily detected by virtue of the highly familiar spatial relationships amongst individual parts (e.g. eyes above the nose and the mouth). When inverted, these relations are disrupted, and recognition can only be achieved through part-based processing, leading to a performance cost in behaviour. Larger inversion effects for social stimuli, including facing dyads, compared to other objects suggest that the visual system may feature specialized perceptual processes that capitalize on the typical configuration in which these stimuli are experienced.

Indeed, just as the face inversion effect is thought to be a marker of the holistic processing of faces (Maurer et al., 2002; Rezlescu et al., 2017; see also Gerlach & Mogensen, 2023), the increased inversion effect elicited by facing *versus* non-facing dyads (Abassi & Papeo, 2022; Papeo, 2020; Papeo et al., 2017) may indicate that facing dyads – like faces – are processed holistically. This has a few important implications. First, it suggests that individuals in facing dyads are bound together perceptually and processed as a unit, in a manner that is more similar to the processing of single bodies or faces than to independent perceptual objects. Second, it suggests that humans have a visual specialisation for facing dyads, which may function as a basic ‘template’ that perceptually highlights social interactions. Importantly, even ‘simple’ facing dyads like the ones used in the current study, isolated from context and background, are viewed as a meaningful social scene (Paparella & Papeo, 2022), and these ‘scenes’ are judged to contain more meaning, emotionality, and intentionality when dyads face each other than when they face away (Goupil et al, 2023). Indeed, even infants and toddlers expect facing (but not facing-away) dyads to interact with each other (Augusti et al, 2010, Beier & Spelke, 2012; Thorgrimsson et al, 2015). Dyadic facing direction, then, is a powerful cue to social interaction – one that could serve as an excellent ‘first-pass’ predictor of when a social interaction is taking place in an observed scene. Visual specialisation for the specific configuration formed by facing dyads may allow for the quick detection of social interactions in the world. Thus, the two-body inversion effect (from here, “2BIE”: the stronger inversion effect for facing vs. non-facing bodies) may be a behavioural marker for the first stage of social interaction perception and processing. One advantage of such a basic perceptual pattern is that this would facilitate attention to social interactions, supporting more efficient cognitive analysis of the observed interaction.

What are the brain mechanisms supporting this proposed visual specialisation for facing dyads? Recent studies have generated new neuroscientific evidence about the perception of interacting dyads. Studies focused on *dynamic* interactions (Isik et al., 2017; Lee Masson & Isik, 2021; Walbrin et al., 2018) have highlighted a region of the posterior superior temporal sulcus (pSTS) as being selectively involved in the perception of observed social interactions, including interactions depicted by full-cue movies (e.g. Landsiedel et al., 2022) animated shapes (Castelli et al., 2000; Walbrin et al., 2018), point-light animations (Centelles et al., 2011; Sapey-Triomphe et al., 2017; Walbrin et al., 2018), and animated mannikins (Georgescu et al., 2014). This region, however, does not respond to *static* depictions of interactions (Landsiedel et al., 2022), including the kind of “prototypical” facing dyads that so reliably produce the 2BIE. As such, it is unlikely that the pSTS is centrally involved in the early stages of social interaction detection described above.

Indeed, other recent neuroimaging evidence implicates more posterior occipitotemporal regions in the perception of interacting dyads, even in a static format. In particular, several studies have suggested the relevance of the “extrastriate body area” (EBA; Downing et al., 2001) in perception of both static and dynamic dyads. To date, most of the research on the functional properties (Downing & Peelen, 2011) and causal relevance (Downing & Peelen, 2016) of this region for social vision has focused on the perception of individual bodies or parts of the body. But recent studies using static images of dyads who are either facing each other as if interacting, or not, have suggested that the EBA is uniquely engaged by facing human dyads (Abassi & Papeo, 2020, 2022). Importantly, work investigating which regions of the brain are most involved in the detection of facing dyads, and the two-body inversion effect, suggests that the size of the 2BIE is directly and specifically related to activity in the EBA (Abassi & Papeo, 2022).

To date, however, evidence for the contribution of EBA to interaction perception remains correlational and limited to single fMRI studies. Important causal evidence about the functional roles of regions in the lateral occipitotemporal cortex (LOTC) in visual processing of single items such as faces, bodies, or objects has come from transcranial magnetic stimulation (TMS) studies (Pitcher et al., 2009). With online TMS, for example, the contribution of category selective regions including the occipital face area (Pitcher et al., 2007), the lateral occipital complex (Dilks et al., 2013; Wischnewski & Peelen, 2021a; Wischnewski & Peelen, 2021b), and EBA (Gandolfo & Downing, 2019; Urgesi et al., 2004; van Koningsbruggen et al., 2013), has been explored. These studies show that TMS selectively impairs performance of tasks requiring attention to images of people or objects in a region- and task-specific manner (Pitcher et al., 2008). Accordingly, they suggest the possibility that a similar interference approach may test hypotheses about the causal contribution of targeted brain areas to interaction perception.

In the current study (see **Figure 1**), we first provide solid correlational evidence for a contribution of the occipito-temporal cortex in interaction perception via a re-analysis of a large neuroimaging dataset. Consistently across participants, results show strong selectivity in the left EBA for visual perception of facing dyads. We then demonstrate the specific and causal role of this region in such functional specialization, using fMRI-guided brain stimulation. In particular, we applied online TMS to the left EBA (localised individually with fMRI) while participants performed a categorization task to measure the 2BIE. The typical pattern of performance in this task, taken as a marker of the visual functional specialization for facing bodies (Papeo et al., 2017; Papeo & Abassi, 2019), is that the effect of inversion on object recognition is more severe for facing than for non-facing dyads, and this does not generalise to control non-body objects (chairs). We replicated the 2BIE in a new sample of participants, and then showed that this facing-direction and category-specific inversion effect was effectively eliminated with online application of TMS over left EBA. In sum, with this rich set of results, we highlight a previously uncharted visual processing stage in social interaction perception and connect it mechanistically to functional specialization in the human visual cortex.

**Figure 1.**
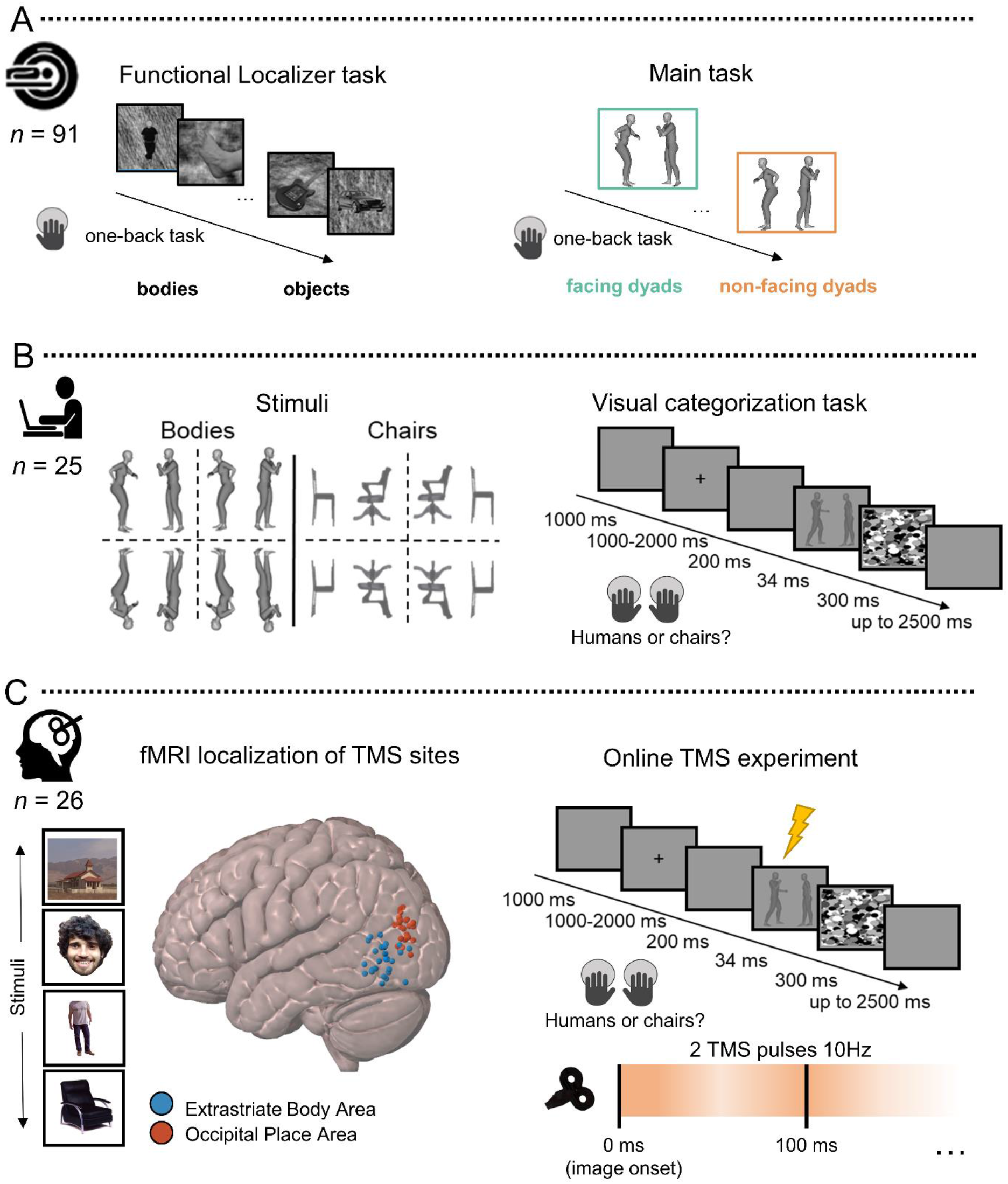
Workflow and schematic display of the procedure across the three experiments. The multi-method approach includes correlational and causal neuroscientific methods. As such, it allows us to directly link brain activity in focal regions with behavioural outcomes. **A.** fMRI study. Left: Examples of stimuli used in the functional localiser task. Extrastriate body area (EBA) was localised with the contrast: bodies > [faces, places, & objects]; Occipital place area (OPA) was localised with the contrast places > [faces, bodies, & objects]. Right: Example of stimuli for key fMRI contrast: facing > non-facing dyads; main experimental task. **B.** Behaviour-only experiment. Left: Examples of experimental stimuli. On each trial, a pair of bodies or chairs were shown upright or inverted, and either facing toward or away from each other. Right: Visual categorization task structure and trial procedure to measure the two-body inversion effect (2BIE). **C.** fMRI-localisation guided TMS experiment. Left: stimuli from the localiser task (the face illustrated was not presented in the real experiment and is the face of the Author (MG)). The localiser session outcomes are displayed on a 3D MNI brain template as normalised individual stimulation site coordinates for the EBA (in blue) and the OPA (in red) for each participant. Bottom: the rTMS session procedure and parameters.

## Results

### Interacting human dyads preferentially engage the left EBA: fMRI evidence

We first performed a new whole-brain fMRI analysis by combining data from 92 participants in three independent fMRI studies (Abassi & Papeo, 2020, 2022, 2023). In each of these studies, participants saw the same critical conditions (facing and non-facing body dyads, **Figure 1A**). While the response in bilateral EBA to facing vs. non-facing dyads was reported in these studies, this reanalysis, done in a properly large sample to provide adequate power, was crucial to our ability to assess the relative response in left vs. right EBA. We used a random-effects general linear model (RFX GLM) to estimate the effect of the facing direction between bodies in a dyad with the contrast facing > non-facing body dyads. Using a voxelwise threshold of p ≤ 0.05, corrected for multiple comparisons using familywise error (FWE) at the voxel level, this contrast (**Figure 2A**) showed an effect in a bilateral cluster within the LOTC peaking in the middle occipital gyrus for the left hemisphere and in the middle temporal gyrus for the right hemisphere. Comparison with the results of a blocked-design functional localizer experiment revealed considerable overlap of this cluster with the EBA at the group-average level. The cluster in the left hemisphere was larger than the cluster in the right hemisphere (Left: 354 voxels, peak MNI coordinates: -52, -76, 12, peak z = 6.26; Right: 14 voxels, peak MNI coordinates: 48, -66, 6, peak z = 4.97, **Figure 2A**). To assess for a significant difference of the overall level of activity between these two clusters, we performed regions of interest (ROI) analyses as follows.

**Figure 2.**
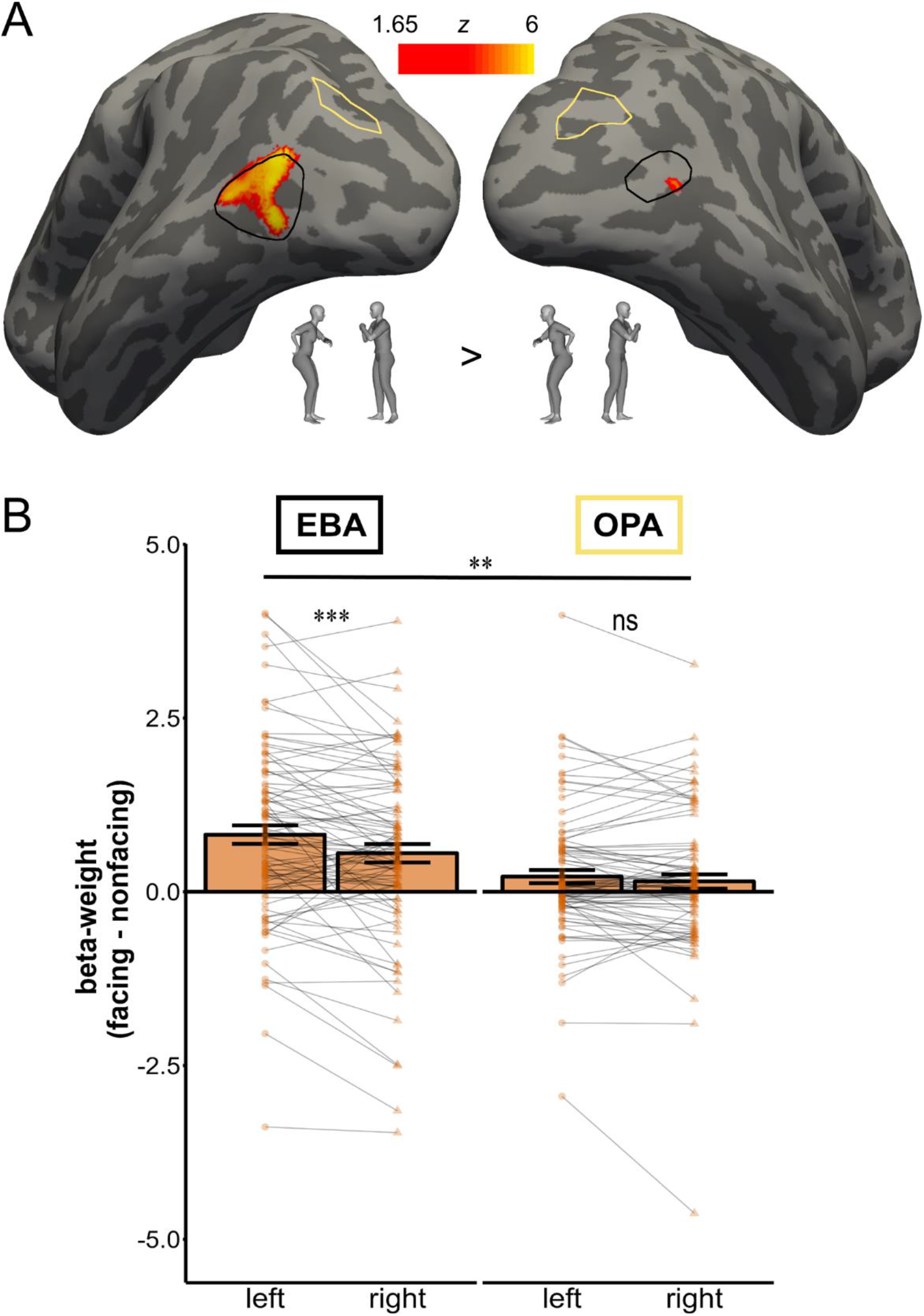
Comparison of brain activity between facing and non-facing dyads. **A.** Univariate second-level (group) analysis for the whole-brain contrast facing > non-facing (N = 92). Voxelwise threshold of p ≤ 0.05, FWE corrected at the voxel level. The EBA highlighted in black corresponds to the results of the group-level random-effect contrasts of bodies > [objects+faces+places]. The OPA highlighted in yellow corresponds to the results of the group-level random-effect contrast of places > [objects+faces+bodies]. **B.** ROI analysis. The bar chart shows the facing > non-facing effect in the EBA and OPA for the left and the right hemisphere. Left hemispheric activity shows a significantly larger difference between the two conditions in the EBA only.

Using data from the functional localizer task, for each participant we identified two ROIs, separately for each hemisphere: the EBA by taking the 200 voxels with the highest selectivity for body-stimuli within an anatomical mask of the middle occipitotemporal cortex, and the OPA by taking the 200 voxels with the highest selectivity for scene-stimuli within the same anatomical mask. To measure the lateralization of the effect of viewing a facing dyad as compared to a non-facing dyad, for each ROI we directly compared the mean neural activity over participants, for facing and non-facing dyads within each hemisphere. A repeated-measures ANOVA with three factors, ROI (EBA/OPA), hemisphere (Left/Right), and facing direction (facing / non-facing), showed a significant three-way interaction (F(1,91) = 7.36; *p* = 0.008; η_p_^2^ = 0.07, **Figure 2B**). Within the EBA-ROI, we found a significant hemisphere by facing direction interaction (F(1,91) = 13.71; *p* < 0.001; η_p_^2^ = 0.13). There was a larger difference in neural response between facing and non-facing dyads in the left EBA (t(91) = 6.20, *p* < 0.001 , d = 0.65, Mean difference = 0.82, 95% CI [0.44, 0.85], BF_10_ = 696000), than in the right EBA (t(91) = 4.19, *p* < 0.001 , d = 0.44, Mean difference = 0.55, 95% CI [0.23, 0.64], BF_10_ = 281, see **Figure 2B**). In contrast, in our control OPA-ROI, we did not observe a significant hemisphere by facing direction interaction (F(1,91) = 2.41; *p* = 0.124; η_p_^2^ = 0.03, BF_10_ = 0.37). These findings together confirm that the facing direction effect was specific to the EBA, though sensitivity to facing direction may also be seen in the fusiform body area (FBA; see supplemental materials), and preferentially lateralized to the left hemisphere.

### The behavioural two body inversion effect (2BIE)

Our next step was to test the hypothesis that the left EBA is causally necessary for the processing of facing human dyads (**Figure 3**). To do so, we first replicated the 2BIE (Papeo et al., 2017). Participants performed a categorisation task to discriminate briefly-presented images of people *vs* chairs (**Figure 1B**). Each image contained two items (people or chairs) that were facing toward or away from each other, and orthogonally either upright or inverted. The 2BIE occurs when the facing human dyads show larger inversion effects – measured by an increase in errors -- than their non-facing counterpart (**Figure 3**). In contrast, in line with prior findings, we did not expect to see similar effects for the control objects (chairs). In line with prior work, our primary measure was accuracy, which is reported in the main text. Reaction time and inverse efficiency score analyses are congruent with the results reported using accuracy in the main text and can be found in the supplementary materials (see **Figure S3**) for both behavioural and TMS results.

**Figure 3.**
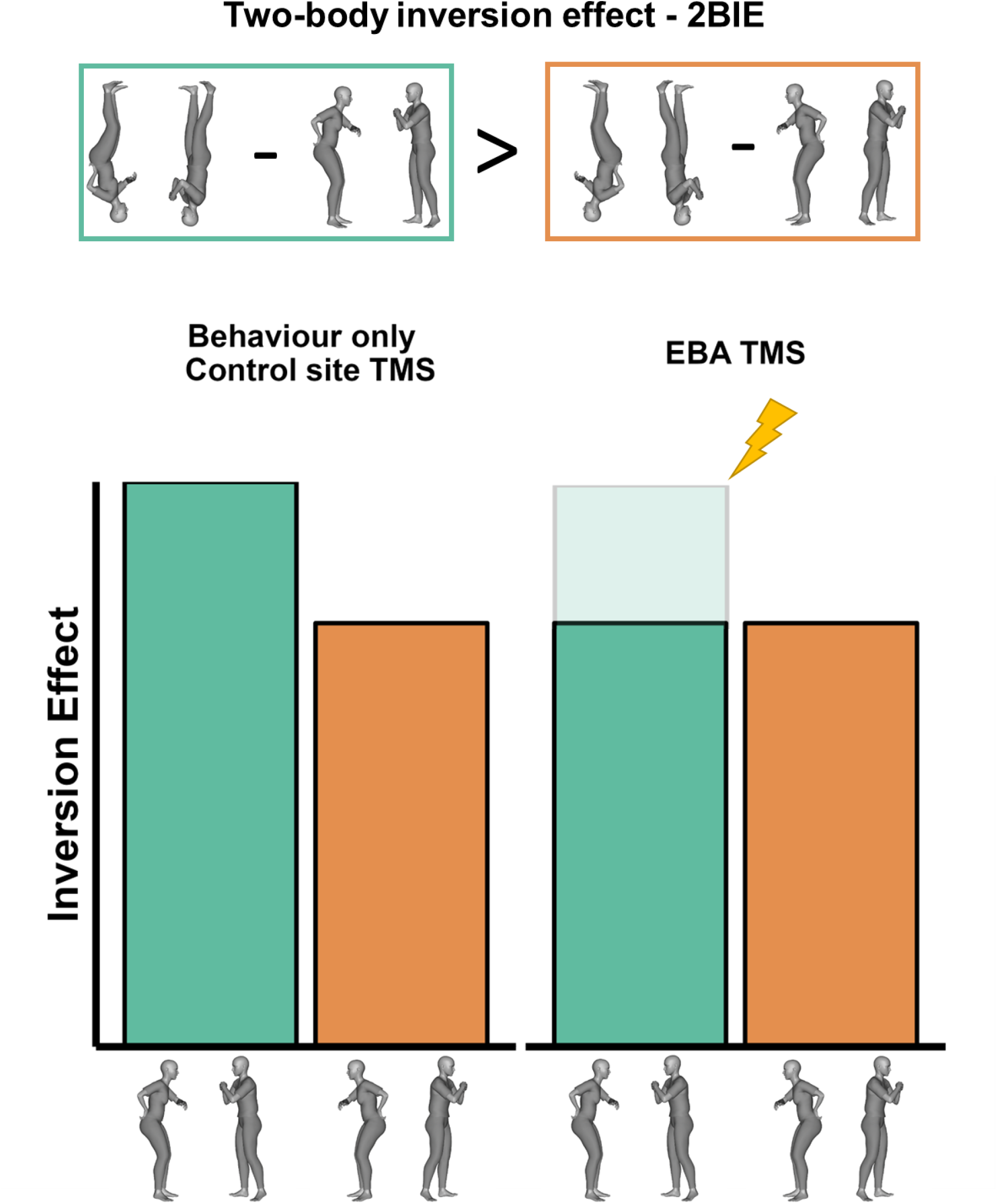
The 2BIE (two-body inversion effect) is taken as a behavioural marker for first-stage social interaction processing. This is measured as increased inversion effects (i.e. difference in performance between inverted and upright body categorization) for facing vs. non-facing dyads. Here, we show the expected pattern of results on the 2BIE for the fMRI guided TMS experiment. EBA stimulation is expected to selectively disrupt the 2BIE while effect should remain intact (i.e., similar to the 2BIE measured in a behaviour only task) after stimulation of the control site (i.e. the OPA).

As predicted, we observed a greater inversion effect for facing dyads than for non-facing dyads, as revealed by a significant three-way interaction among Stimulus (bodies, chairs), Orientation (upright, inverted), and Facing direction (facing, non-facing), F(1,22) = 7.43, *p* = 0.012, η_p_^2^ = 0.25. For human stimuli, the inversion effect for facing dyads was larger than for the non-facing dyads (**Figure 4A**, t(22) = 3.014, *p =* 0.006, d = 0.63, Mean Difference = 6.93%, 95% CI [2.16, 11.70], BF_10_ = 7.16). This difference was absent when comparing the inversion effect for facing and non-facing chairs (**Figure S2**) (t(22) = -0.51, *p* = 0.616, d = - 1.22, Mean Difference = 1.22%, 95% CI [-6.21, 3.76], BF_10_ = 0.25; See supplemental materials). We take this facing-direction and category-specific inversion effect as the behavioural marker of early-stage visual processing of social interaction that we then targeted in the subsequent neurostimulation study.

**Figure 4.**
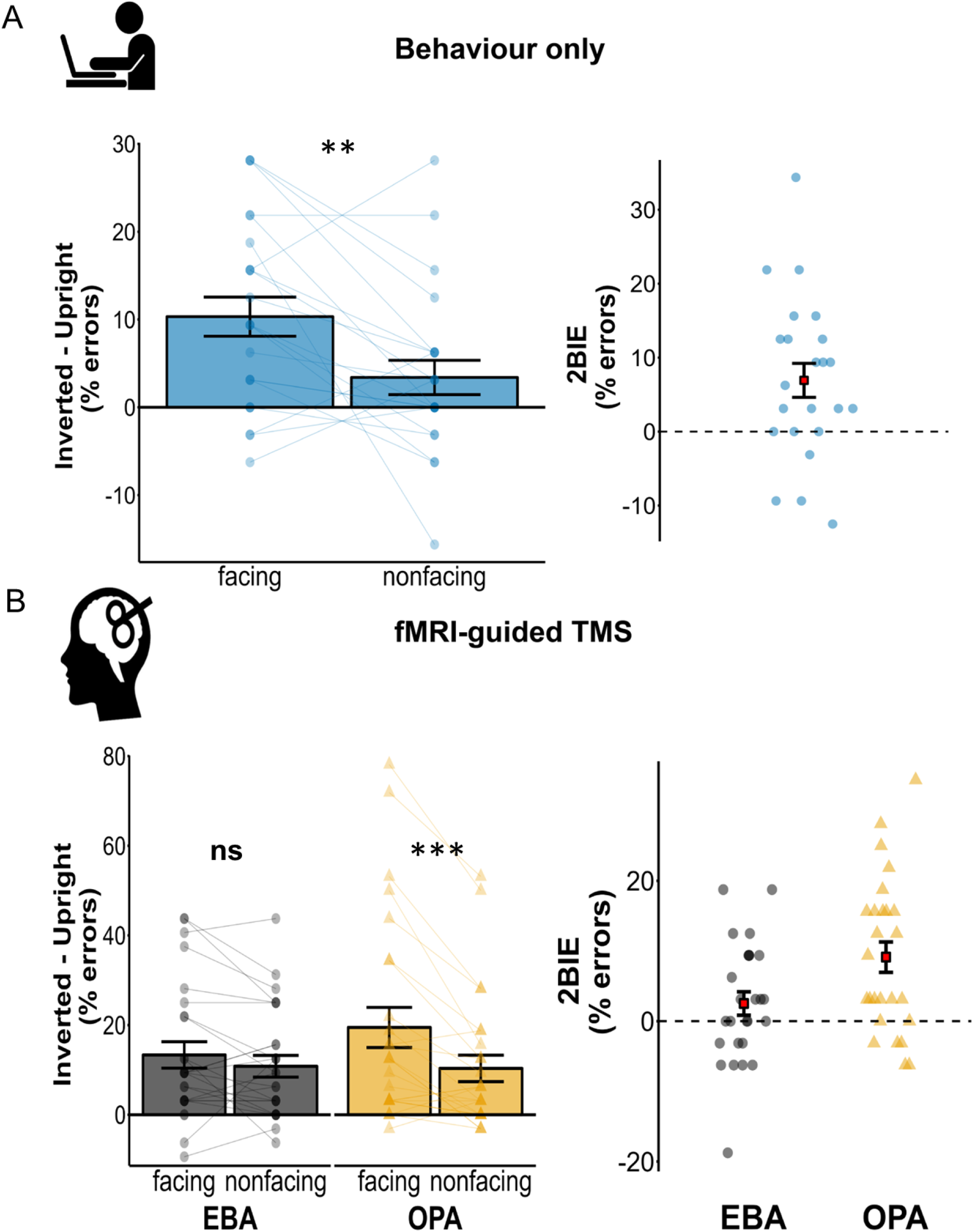
**A.** Behaviour-only experiment (N = 25). The left panel bar chart shows the inversion effect (% of errors in the inverted – upright trials) for human dyad stimuli in the facing vs non-facing condition. The right panel illustrates the 2BIE, expressed as the subtraction of the inversion effect for non-facing dyads from that for facing dyads. **B.** fMRI guided TMS-experiment (N = 26). The left panel shows the inversion effect (% of errors in the inverted – upright trials) for human dyad stimuli in the facing vs non-facing condition for both the EBA (site of interest) and the OPA (control TMS site). The right panel shows the 2BIE (inversion effect for facing – inversion effect for non-facing dyads) during EBA vs OPA stimulation. TMS on the EBA reduced the 2BIE; TMS on the OPA (control site) yielded a pattern of results consistent with that seen in the behaviour-only experiment (note that the behaviour-only and fMRI-guided TMS experiment were conducted with different participants). The full barchart including all of the condition means can be found in **Figure S1.** Lines and datapoints indicate individual participant’s means in the respective conditions. Error bars indicate the standard error of the mean. * p < .05; ** p < .01; *** p < .005.

### Left-EBA is causally necessary for encoding facing human dyads

If activity of the left hemisphere EBA directly contributes to early-stage processing of static social interactions, as indicated by our correlational evidence from fMRI, then online stimulation of this region should selectively disrupt the 2BIE (**Figure 3**). Further, this TMS effect should be category-specific (only occurring with human dyads) and site-specific (only occurring after left EBA stimulation). Note that these hypotheses were pre-registered (https://aspredicted.org/4CZ_GGG).

As predicted, we found that left EBA stimulation selectively eliminated the 2BIE, and therefore the TMS effects on the categorization task were category- and site-specific (**Figure 4B**). This was revealed by a significant site (EBA, OPA) x stimulus (body, chair) x orientation (upright, inverted) x facing-direction (facing, non-facing) interaction, F(1,25) = 4.92, *p* = 0.036, η_p_^2^ = 0.16.

To clarify the nature of this interaction, we conducted separate three-way ANOVAs for each region of interest. When TMS was applied to the control site OPA, the selective effect of inversion on facing dyads was confirmed, similar to the 2BIE found without TMS (behaviour-only experiment), as evidenced by a Stimulus x Orientation x Facing-direction interaction (F (1,25) = 9.90, *p* = 0.004, η_p_^2^ = 0.28, BF_10_ = 9.79). In contrast, when TMS was applied to EBA the interaction that characterizes the 2BIE was not significant (F (1,25) = 0.31, *p* = 0.581, η_p_^2^ = 0.01). Further, a Bayesian analysis of this contrast indicated positive evidence for the null hypothesis (BF_10_ = 0.24).

Indeed, when directly comparing the effect of stimulation at the two stimulation sites when participants were viewing dyads of people, we found a site x orientation x facing-direction interaction (F(1,25) = 9.87, *p* = 0.004, η_p_^2^ = 0.28). That is, the inversion effect for the facing human dyads was larger than for their non-facing counterpart after left OPA stimulation (t(25) = 4.22, *p <* 0.001, d = 0.83, Mean Difference = 9.13%, 95% CI [4.68, 13.59], BF_10_ = 105.79). In contrast, those conditions did not reliably differ after left EBA stimulation (t(25) = 1.51, *p =* 0.144, d = 0.30, Mean Difference = 2.52%, 95% CI [-0.92, 5.97], BF_10_ = 0.57). Additionally, a paired t-contrast on the inversion effects in EBA vs OPA for the facing human dyads confirmed that EBA stimulation affected the inversion effects in the facing condition (t(25) = 2.29, p = 0.030, d = 0.45, Mean Difference = 6.13%, 95% CI = [0.63, 11.63], BF_10_ = 1.88), while the same contrast did not reach significance in the non-facing condition (t (25) = -0.22, p = 0.830, d = -0.04, Mean Difference = 0.48%, 95% CI = [-5.05, 4.09], BF_10_ = 0.21).

Finally, no difference between facing and non-facing orientation was found for the control chair stimuli (F(1,25) = 0.62, *p* = 0.438, η_p_^2^ = 0.02, **Figure S2**) demonstrating that neither left EBA (t(25) = 0.39, *p =* 0.702, d = 0.08, Mean Difference = 0.96%, 95% CI [-4.15, 6.07], BF_10_ = 0.22) nor left OPA (t(25) = -0.86, *p =* 0.397, d = 0.17, Mean Difference = 2.52%, 95% CI [-0.57, 0.24], BF_10_ = 0.29) stimulation affected performance in this condition. Together, these analyses confirm that the effects of TMS on the 2BIE were both site and category specific, as predicted.

## General Discussion

Here, we present the first causal evidence for the contribution of occipitotemporal social perception regions, specifically left EBA, to the perception of interacting (face-to-face) human dyads. First, we demonstrate across a large group of participants that left EBA, in particular, is more responsive to facing dyads than non-facing dyads. We then demonstrate that TMS to left EBA effectively eliminates the two-body inversion effect, which is proposed to constitute a signature of early processes in social interaction perception. The specificity of this result to human stimuli (*vs* chairs) demonstrates that it is not due to a generic disruption of visual processing. The specificity of this result to left EBA stimulation (*vs* nearby OPA) demonstrates that reduction of the 2BIE does not necessarily follow disruption of any given occipitotemporal region, nor is it due to non-specific ‘annoyance’ caused by TMS, such as task interference due to stimulator noise, or discomfort caused by TMS pulses (see Meteyard & Holmes, 2018).

What is the nature of the causal contribution made by left EBA to perceiving dyadic interactions? It is possible that disrupting EBA’s encoding of individual body postures partly contribute to the TMS effects reported here. Specifically, TMS to EBA may reduce the facing-specific inversion effect by disrupting body-posture signals that would normally help distinguish facing from non-facing pairs. This contribution of EBA may be particularly strong for inverted postures: if inversion disrupts whole-body representations, then encoding of the body posture must rely more on part-based body representations. Indeed, previous evidence from fMRI (Taylor et al., 2007; Taylor & Downing, 2011; Urgesi et al., 2007) indicates an emphasis on body-part representations in EBA, relative, for example, to whole-body representations in the fusiform body area (FBA; Peelen & Downing, 2005; Schwarzlose et al., 2005). This account, however, cannot fully explain our results because it also predicts a general disruption of the two-body inversion effect regardless of facing direction – a pattern that is not seen in our data. Likewise, it does not easily account for the increased response to facing over non-facing dyads in left EBA in our, and others, fMRI data.

Instead, we suggest that TMS to left-EBA disrupts the holistic processing of facing dyads. Indeed, we used the two-body inversion effect as a behavioural marker of early interaction processing because it is thought to relate to holistic processing of prototypical human face-to-face interactions when dyads are seen in their canonical upright orientation (Papeo et al., 2017). Thus, left EBA activity may normally capture an emergent property of facing dyads that reflects their proposed status as a “unit” of visual processing (Adibpour et al., 2021), beyond its response to individual bodies. This idea is in line with previously observed sensitivity for other kinds of typically-observed object configurations in the broader LOTC (Kaiser et al., 2019; Kaiser & Peelen, 2018; Papeo, 2020). Importantly, if left EBA is involved in holistic processing of facing dyads, this both fully explains our TMS results and accounts for left EBA’s specific sensitivity to facing over non-facing dyads in our fMRI results.

Our demonstration of a relatively greater fMRI response to static facing dyads in left rather than right hemisphere EBA contrasts with a general tendency for the effects of social stimuli (people and their movements) on visual cortex activity to be relatively right-hemisphere lateralised (e.g., Allison et al., 1999; Downing et al., 2001; Isik et al., 2017; Kanwisher et al., 1997). Indeed, as a result of this right-lateralisation, much of the literature investigating responses in social perceptual regions (e.g., FFA, EBA) has investigated *only* regions in the right hemisphere. More generally, evidence from neuropsychological studies suggests a relatively greater involvement of the right hemisphere in holistic or global perception and, in contrast, an emphasis on parts or local processes in the left hemisphere (e.g., Robertson, 1995). How to reconcile our findings with these established patterns? One intuitive possibility is that our result could be related to the left-lateralised hand-selective region that is near to, but distinct from, EBA (e.g., Bracci et al., 2010). When we observe dyadic interactions in the world, people are more likely to be reaching towards than away from each other. When presenting those stimuli in the lab, extremities (hands, feet) are thus more likely to appear on or near the fovea in facing than in non-facing dyads, which could drive hand-selective regions more strongly for facing dyads. This is not, however, the case for our stimuli, where the distance between the nearest point (usually hands or feet) and the screen centre was matched between facing and non-facing dyads, across stimuli. In addition, the distance between hands and/or feet is *exactly* the same for inverted dyads as for upright dyads. Thus, a strong visual representation of the hands in the left hemisphere would not in itself account for the present fMRI or TMS results. Instead, we propose a related possibility, that sensitivity to social interactions in left EBA is related to findings that many aspects of actions are represented in the wider lateral occipitotemporal cortex, and particularly so in the left hemisphere (Lingnau & Downing, 2015). Our fMRI results may reflect the increased processing needed to represent body and limb postures that predict interaction-specific actions when dyads face each other. Thus, our results are not necessarily in conflict with what we already know about EBA, but instead may show a complementary functional specialisation in left EBA for representing body information that is specifically relevant to predicting joint, coordinated, or reciprocal (interactive) actions.

Here, we have focused on the role of EBA in interaction perception. The fusiform body area (FBA) also shows some sensitivity to cues to dyadic interactions (Supplemental materials; Abassi & Papeo, 2020, 2023) yet the causal role for this ventral region on the 2BIE could not be directly assessed as it is not reachable by TMS. Additionally, previous evidence suggests that the fMRI sensitivity for facing dyads measured in the FBA, unlike the EBA, is not reliably related to the 2BIE (Abassi & Papeo, 2022). Similarly, prior work suggests that bilateral pSTS is also strongly sensitive to visual cues to dyadic interaction, including facing direction. However, pSTS appears to only be sensitive to such interactive cues when they are presented dynamically (Landsiedel et al., 2022), while EBA shows sensitivity to dyadic facing direction for both static and dynamic stimuli (Abassi & Papeo, 2020; Bellot et al., 2021; Landsiedel et al., 2022; Walbrin & Koldewyn, 2019). Additionally, at least for dynamic social stimuli, visual information is proposed to feed from EBA to pSTS (Giese & Poggio, 2003; Pitcher & Ungerleider, 2021). Indeed, recent evidence shows that functional connectivity between EBA and pSTS is increased during the perception of dyads that move towards vs away from each other (Bellot et al., 2021). Thus, the two regions likely work together during ‘real world’ perception of social interactions, which are necessarily dynamic and unfold across time. This proposal prompts a wealth of future research questions. Is motion necessary for information flow between EBA and pSTS? What, specifically, does pSTS add to the processing of social interactions? How does information flow beyond EBA and pSTS to support higher-level social cognitive processes, social inferences, and social judgements? Our results also specifically prompt a reassessment of the lateralisation within the ‘social brain’, as it will be important to understand the flow of social information between left EBA and right-lateralised social perception regions, particularly right pSTS.

## Concluding remarks

Using a multimodal approach, we demonstrate that left EBA is sensitive to interactive information. Further, we demonstrate that activity of this region is causally necessary for the efficient perception of facing dyads. Rapid detection of facing dyads may reflect a processing stage that is critical for recognising and understanding social interactions in a cluttered visual world. As such, our results strongly support the idea that the first steps towards understanding social interactions are underpinned by basic visual processes, critically relying on discriminating between interacting and non-interacting human groups. Importantly, although future work is needed to fully explore the role EBA plays in social interaction perception and social inference, our results also demonstrate a novel functional specialisation in left EBA.

## Methods

### RESOURCE AVAILABILITY

#### Lead contact

Further information and requests for resources, clarification about the data and code should be directed to and will be fulfilled by the lead contact, Marco Gandolfo ( –).

#### Materials availability

All stimuli are freely available from the authors (write to). A sample of these stimuli are available at the Open Science Framework (OSF) project page associated with this study (OSF | Left EBA stimulation disrupts configural processing of face-to-face human dyads). DOI is listed on the key resource table.

#### Data and code availability

Behavioural, beta values for the ROIs, and TMS data have been deposited at the OSF repository (https://osf.io/m9vp2/). Analyses R scripts have been deposited at the OSF repository (https://osf.io/m9vp2/) and listed on the key resource table. Any additional information required to reanalyse the data reported in this paper is available from the lead contact upon request.

### EXPERIMENTAL MODEL AND SUBJECT DETAILS

#### Participants: fMRI

We combined data from 92 unique participants (52 females and 40 males, self-reported sex; mean age 24.8 years, SD = 3.7) included in one of three previous independent fMRI studies (*N* 20 from Abassi and Papeo, 2020; *N* 30 from Abassi & Papeo, 2022; *N* 42 from Abassi & Papeo, 2023). All participants had normal or corrected-to-normal vision, reported no history of psychiatric or neurological disorders, or use of psychoactive medications. They were screened for contraindications to fMRI and gave written informed consent. All studies were approved by the local ethics committee (CPP Sud Est V, CHU de Grenoble).

#### Participants: Behaviour-only experiment

With this study, we sought to replicate the two-body inversion effect (2BIE) using stimuli, task and procedures from the original study that first reported that effect (Papeo et al., 2017). To this end, 25 participants were recruited (18 females, 7 males, mean age 20, SE = 1.3). This sample size matched Papeo et al. (2017). Additionally, a sensitivity analysis showed that a sample of this size allows to achieve 80% power with an effect size of at least d = 0.58 on a two-tailed t-contrast. Two participants were excluded because their mean accuracy (1 participant) or response times (1 participant) were >2.5 above or below the group mean across conditions. Thus, a final sample of 23 participants were included in the final analyses. For this and the following experiment the procedures were approved by the Research Ethics Committee of Bangor University’s School of Psychology. Participants were community members of Bangor University and provided informed consent before participation. They were compensated with course credits, or a cash payment. Each participant only took part in one experiment.

#### Participants: TMS experiment

All procedures and the following analyses were pre-registered on aspredicted.org (https://aspredicted.org/4CZ_GGG). Deviations from the pre-registrations are indicated in the corresponding sections. Twenty-nine participants (16 females, 12 males, 1 other, mean age 24.34) took part in the TMS study. They were screened following the safety screening standard questionnaire for rTMS (Rossi et al., 2021). None of them reported a history of neurological, psychiatric, or other major medical disorders. Three participants were excluded from further analysis because their accuracy was 2.5 *SD*s below the group mean across conditions, yielding a final sample of 26. A sample size of 25 participants was pre-registered, to match the planned sample size of the above behaviour-only study, where we successfully replicated the 2BIE (see Results). However, to allow full counterbalancing of the stimulation site (participant starting with EBA/OPA stimulation) we tested one additional participant.

### METHOD DETAILS

#### fMRI re-analysis

##### Stimuli

In all three studies, stimuli were created from grayscale renderings of human bodies in various biomechanically possible poses, in a profile view, arranged in pairs, to form face-to-face (facing) dyads or back-to-back (non-facing) dyads. Non-facing body dyads were created by swapping the position of the two bodies in each facing body dyad. Distance between the two bodies in a dyad was matched across facing and non-facign dyads so that the two sets only differed for the relative positioning of the bodies. We used Daz3D (Daz Productions, Salt Lake City) and the Image Processing Toolbox of MATLAB (The MathWorks Inc, Natick, Massachusetts) to create and edit stimuli. For a full description of the stimuli, we refer the reader to Abassi & Papeo (2020; 2022; 2023).

Inside the scanner, a liquid crystal projector (frame rate: 60 Hz; screen resolution: 1024 × 768 pixels, screen size: 40 x 30 cm) was used to back-project the stimuli onto a screen. Participants viewed the stimuli binocularly (∼7° of visual angle) through a mirror above the head coil. Stimulus presentation, response collection, and synchronization with the scanner were controlled with Psychtoolbox (Brainard, 1997) through MATLAB (MathWorks).

##### fMRI data acquisition

Imaging in all three studies was conducted on the same MRI machine, a MAGNETOM Prisma 3T scanner (Siemens Healthcare) at the CERMEP imaging centre in Bron (France). T2*-weighted functional volumes were acquired using a gradient-echo echo-planar imaging sequence with slightly different values across studies due to the fact in studies 2 and 3 multiband acquisitions were employed (Study 1: GRE-EPI; TR/TE = 2200/30 ms, flip angle = 90°, acquisition matrix = 74x74, 40 slices, FOV= 220x220 mm, 40 slices, slice thickness = 3 mm, no gap, acceleration factor = 2 with GRAPPA reconstruction and phase encoding set to anterior/posterior direction; Study 2-3: GRE-EPI; TR/TE = 2000/30 ms, flip angle = 80°, acquisition matrix = 96x92, FOV = 210x201 mm, 56 slices, slice thickness = 2.2mm, no gap, multiband acceleration factor = 2 and phase encoding set to anterior/posterior direction). For all studies, acquisition of high-resolution T1-weighted anatomical images was performed in the middle of the main experiment and lasted 8 min (MPRAGE; TR/TE/TI = 3000/3.7/1100 ms, flip angle = 8°, acquisition matrix = 320x280, FOV = 256x224 mm, slice thickness = 0.8mm, 224 sagittal slices, GRAPPA accelerator factor = 2). Acquisition of two field maps was performed at the beginning of each fMRI session.

##### fMRI task procedure

All studies included three runs featuring blocks of upright facing or non-facing dyads (randomized with other type of runs that varied across study). The structure of blocks was identical across studies, although with slight variation in the duration of events. Each block included 5-to-8 stimuli of the same condition (facing or non-facing) presented for 400-to-550 ms, interleaved with a black central fixation cross, which changed colour (from black to red) in 37.5% of all stimulation and fixation trials. Participants were instructed to fixate the cross throughout the experiment and detect and report the colour change by pressing a button with their right index finger. This task was used to minimize eye movements and maintain vigilance in the scanner. For more information on the specificities of each study, we refer the reader to Abassi & Papeo (2020; 2022; 2023).

##### Functional localizer tasks

For all studies, in addition to the experimental runs, subjects completed a functional localizer task of 2-to-3 runs, with stimuli and task adapted from the fLoc package (Stigliani et al., 2015). In all three studies, the functional localizer task included grayscale photographs of: 1) bodies; 2) faces; 3) places; and 4) inanimate objects. Within a run, the duration of each block was four seconds, with 12 blocks for each object class over the run and eight images per block (500 ms per image without interruption), randomly interleaved with 12 baseline blocks (empty screen).

#### Behavioural and TMS experiments

##### Stimuli

Stimuli included facing and non-facing human body dyads, pairs of facing and non-facing chairs, and visual masks, taken from Papeo et al. (2017). For each category, facing and non-facing stimuli involved the very same items, differing only for the relative positioning. Importantly, the distance between bodies in facing and non-facing dyads was matched by keeping the centre-centre distance of each pair equally distant from the centre of the display (1.8° of visual angle). Moreover, the distance between the two closest points of the two bodies was comparable for facing and non-facing dyads (mean distances for facing in pixels: 42.5 +/- 9.8 SD; mean distances for nonfacing in pixels: 40.3 +/- 12.2 SD; Mann-Whitney *U* test of the difference: *z* = 0.44, *p* = 0.660). A similar approach was implemented for creating pairs of chairs (60 facing and 60 non-facing), starting from six unique exemplars. Like body dyads, pairs of facing and non-facing chairs included the very same items at a matched distance (∼2 degrees of visual angle), and thus only differed for the relative positioning (see Papeo et al., 2017 for details on stimuli). Visual masks consisted of high-contrast Mondrian arrays (11° x 10°) of grayscale circles of variable diameter (0.4°-1.8°). All stimuli were shown on a grey background. Images were presented on an 18 inch CRT monitor screen (1024x768 px, 60Hz refresh rate). The task was coded using Psychtoolbox (Brainard, 1997) running on GNU Octave (Eaton, 2018) on a Xubuntu Linux machine.

##### Task Procedure

Participants were seated on a chair in front of the monitor at a distance of ∼50 cm from the screen in a dimly lit room. For each trial, they were instructed to report whether they saw humans or chairs. Each trial included the following sequential events: a grey screen (200 ms), a central fixation cross (5 jittered durations ranging between 1 and 2 seconds in intervals of 250 ms), the stimulus (upright/inverted facing/non-facing human dyad or pair of chairs) shown for 34 ms (2 screen refreshes at 60hz), and a visual mask shown for 300 ms. Participants were asked to respond “humans” or “chairs” as accurately and as quickly as possible, by pressing <F> or <J> on the keyboard. The key to response mapping was randomised across participants. Every 32 trials, participants were invited to take a break. Before starting, participants performed a practice block of 32 trials. The first 16 practice trials showed the stimuli for 250ms, so that participants could see the stimuli clearly and become familiar with them. In the second half of the practice block, the images were presented at the same speed as in the experiment (34 ms). Participants were offered the opportunity to perform an additional practice block, if needed. In the TMS version of the experiment, everything was identical to the above, except that the experimenter (MG or EB) applied TMS pulses online, with the TMS coil placed on the participant’s scalp.

##### Design

In the behaviour-only study, participants completed 256 trials divided in eight counterbalanced blocks of 32 trials (4 trials for each stimulus category, orientation, and facing direction presented in random order). During the TMS study, participants completed this task twice, once for each stimulation site. The site of first stimulation was counterbalanced across participants.

##### TMS parameters

A Magstim Rapid^2^ (Magstim; Whitland, UK) with a 70mm figure-eight coil was used for the TMS. Stimulation intensity was set at 65% of the maximum stimulator output. This intensity corresponds to the average 120% of the resting motor threshold of Gandolfo & Downing (2019) using the same TMS equipment, and is similar to previous studies (Pitcher et al., 2007, 2008). Online TMS was delivered at 10Hz (2 pulses, 1 pulse every 100ms, 1 pulse at image onset, and 1 pulse 100ms after image onset) with the handle pointing downward approximately at 45° from the middle sagittal axis of the participants’ head (Candidi et al., 2008; Urgesi et al., 2004, 2007), adjusted to best project the pulse to the identified peak coordinate of each region and kept constant across stimulation sites. Such online stimulation protocol has been widely used to interfere with the ongoing processing of the preferred stimuli of category selective regions in occipito-temporal cortex (Pitcher et al., 2012; van Koningsbruggen et al., 2013).

TMS targeting was monitored with Brainsight 2.3.11 (Rogue Research), using individual structural and functional MRI images for each participant. The left EBA and left OPA were localized by overlaying individual activation maps from the localizer contrasts. The coil location was monitored online by the experimenter (Authors MG or EB) while participants performed the task, and was maintained within 1mm of the defined point. The screen displaying the participants’ task was out of view of the experimenters, rendering them blind to the condition on a trial-by-trial basis. To ensure temporal precision, the train of TMS pulses was triggered on each trial via a TTL pulse, initiated from a photosensor which detected a screen event (unseen by the participant) that co-occurred with the image onset on each trial.

##### Target hemisphere

Our pre-registration anticipated the stimulation of the right EBA, given that visual body-related effects consistently show right lateralisation (Gandolfo & Downing, 2019; Urgesi et al., 2007; van Koningsbruggen et al., 2013; Vangeneugden et al., 2014). However, an analysis on a pilot study involving 12 participants showed no modulation of the 2BIE, or any effect of TMS on task performance (see Supplemental Information). In parallel, we obtained the results of the fMRI re-analysis, which showed a strongly left-lateralized effect of facing *vs.* non-facing body dyads (see Results). Therefore, we reinvited the above 12 participants for another TMS study, in which everything was identical to the previous one except for the stimulation site: left EBA/OPA rather than right EBA/OPA. Eight out of 12 participants returned to the lab for this second study (COVID19-related disruptions caused the remaining participants to leave Bangor University before they could join the experimental session). Thus, both our fMRI results and these preliminary TMS analyses on this subset of participants (See supplemental materials, **Figure S4**) pointed to the left (vs. right) hemisphere as being involved in the current effect. For this reason, and to avoid unnecessary brain stimulation, the remaining participants reported in the TMS study were only stimulated on the left ROIs. All the other procedures remained the same as described in the pre-registration. Data for the twelve participants on the right hemisphere are available on the OSF (OSF | Left EBA stimulation disrupts configural processing of face-to-face human dyads).

### QUANTIFICATION AND STATISTICAL ANALYSIS

#### fMRI reanalysis

##### Preprocessing of fMRI data

Functional images were pre-processed and analysed using SPM12 (Friston et al., 2007) as implemented in MATLAB. The first four volumes of each run were discarded, taking into account initial scanner gradient stabilization (Soares et al., 2016). The pre-processing of the remaining volumes involved de-spiking, slice time correction, geometric distortion correction using field maps, spatial realignment and motion correction using the first volume of each run as reference. Anatomical volumes were co-registered to the mean functional image, segmented into grey matter, white matter and cerebrospinal fluid, and aligned to the Montreal Neurological Institute (MNI) template. To normalise the functional images, we used the DARTEL method (Ashburner, 2007) to create flow fields for each subject and an inter-subject template, registered in the MNI space. Finally, the voxels were resampled to 2mm^3^, spatially smoothed with a Gaussian kernel of 6 mm FWHM and low-frequency drifts were removed with a temporal high-pass filter (cutoff 128 s).

##### Whole-brain univariate analyses

Random-effects general linear model (RFX GLM) analyses were conducted to estimate the blood-oxygen-level-dependent (BOLD) signal. The model comprised two regressors for the experimental conditions (facing and non-facing dyads), one regressor for the fixation blocks, and six nuisance regressors for movement correction parameters. The facing > non-facing effect in the second-level (group) analysis was determined using a voxelwise threshold of p ≤ 0.05 which was family wise error (FWE) corrected at the voxel level to correct for multiple comparisons.

##### Definition of regions of interest (ROIs)

For each subject, using data from the functional localiser task, we identified voxels showing the highest response to bodies in the bilateral EBA using the contrast bodies > faces+objects+places, and to places in the bilateral OPA, using the contrast places > faces+objects+bodies. Individual subject localiser data were entered into a GLM with one regressor for each object-class condition (bodies, places, faces and objects), one for baseline blocks, and six nuisance regressors for movement correction parameters. One mask of the left and one of the right middle occipitotemporal cortex was created using FSLeyes (McCarthy, 2018) and the Harvard-Oxford Atlas (Desikan et al., 2006) implemented in FSL (Jenkinson et al., 2012). For each participant, within this mask, we first selected the voxels with significant activation (threshold: p = 0.05) for each of the above two contrasts. For each individual, the two final ROIs included up to the 200 voxels with the highest t-values.

##### ROIs analyses

The mean activation estimation parameters for facing and non-facing dyads were extracted and analysed in a 2 ROI (EBA/OPA) x 2 hemispheres (left / Right) x 2 facing-direction (facing/non-facing) repeated-measures ANOVA.

#### Behavioural and TMS experiments

Statistical significance was tested with factorial design ANOVAs and follow-up t-tests. Significance level was set at p = 0.05. In line with previous reports with this task we report analyses of Accuracy (% errors), computed for each condition. We performed a 2x2x2 within-subjects ANOVA with Stimulus (People/Chairs) x Orientation (Upright/inverted) x Facing Direction (Facing/non-facing) for the initial behaviour-only study. In the case of a significant interaction we compared the inversion effect (% errors for upright minus inverted stimuli) for facing *vs* non-facing conditions for each stimulus. In the TMS study, the design was the same, with the additional within-subjects factor of stimulation site (EBA/OPA). The interaction among all factors was further inspected with two separate ANOVAs, comparing the effects between the two types of stimuli, separately for each site, as well as two other ANOVAs comparing effects between the two stimulation sites, separately for each stimulus. Alongside the paired t-tests, we report the mean difference in percentage of errors between conditions and the 95% confidence interval of this difference.

Where the absence of the effects was informative for the results, we added Bayes Factors as a complement to the null-hypothesis significance testing. We report BF10 calculated based on an uninformative standard Cauchy prior. Briefly, BF10 below 1 indicate that the null hypothesis is more likely, and values below 1/3 are generally taken as moderate, reliable evidence in favour of the null hypothesis (Jeffreys, 1961; Lee & Wagenmakers, 2013). For visualisation purposes, we calculated indexes subtracting the inversion effect for facing vs non-facing stimuli for each stimulation site. When such measure is positive, it indicates larger inversion effects for facing – vs – non-facing stimuli (Papeo et al., 2017). Data pre-processing and analyses were conducted using R (Version 4.2.1) (R Core Team, 2022).

##### fMRI-based localisation of TMS sites

Each participant in the TMS study first completed two to four runs of a functional localizer task in the fMRI on a different date (∼ 1 week before the TMS session). The task involved four conditions (human bodies without heads, unfamiliar faces, outdoor scenes, and chairs), each presented in four blocks of 18s in each run, interspersed with 5 fixation blocks of 16 s duration, for a total of 21 blocks per run. In each block, 24 images (selected randomly from a full set of 40) were presented, each for 300 ms, followed by a 450 ms blank interval. Twice during each block, an image was presented twice in a row. Participants were instructed to detect and report these repetitions by pressing a key.

Imaging data were acquired using a 3T Philips MRI scanner with a 32-channel SENSE phased-array head coil at the Bangor Imaging Centre in Bangor, Wales (UK). Functional data (T2* weighted, gradient echo sequence, echo time, 35ms; flip angle, 90°) were acquired with the following scanning parameters: repetition time 2s; 35 off-axial slices; voxel dimensions 3x3 mm; 3mm slice thickness; SENSE factor 2, phase encoding direction anterior-posterior. A high-resolution anatomical scan was also acquired (T1 weighted, 175 sagitally oriented slices; 1mm isotropic voxels; repetition time, 8.4 ms; echo time, 3.8 ms; flip angle 8°).

Functional MRI data were preprocessed and analysed using SPM12 (Statistical Parametric mapping software; Wellcome Trust Centre for Neuroimaging, London, UK). The functional images were realigned, co-registered with the anatomical T1 image and spatially smoothed (6-mm FWHM Gaussian kernel). The resulting images were entered into a subject-specific general linear model with four conditions of interest corresponding to the four categories of visual stimuli. Estimates of the BOLD response in each voxel and category were derived by a general linear model including the boxcar functions of stimulation that were convolved with a standard hemodynamic response function. All analyses were performed in participant-native coordinates; for reporting purposes the coordinate of target sites were converted to standard MNI space (see **Figure 1C**).

For each participant, we localized the left hemisphere body and scene selective regions by contrasting the response to human bodies with that to the remaining three conditions and the response to scenes with that to the remaining three conditions, respectively. Each TMS target site (left hemisphere EBA; left OPA) was individually identified by selecting the peak activation for the relevant contrast, in the left lateral occipito-temporal cortex. The mean peak MNI coordinates (X, Y, Z, with SEM) were -45 (0.99), -73 (1.32), -2 (1.71) for the left EBA, and -32 (0.77), -84 (0.62), 17 (1.51) for the left OPA (see **Table S1**, **Figure 1C** for individual coordinates).

## Declaration of Interests

The Authors declare no competing interests.

## Acknowledgments

S. McKiernan for development of the custom photo-diode trigger, Sarah Salzgeber for task development and data collection in the behavioural experiment, Violette Munin for her help in the pre-processing and analysis of the fMRI data, and the Koldewyn lab for helpful discussions during lab meetings.

## Funding

This project has received funding from the European Research Council under the European Union’s Horizon 2020 research and innovation programme (ERC-2016-STG-716974: Becoming Social; awarded to K. Koldewyn), (ERC-2017-STG-758473 – THEMPO: awarded to L. Papeo), and the Marie Skłodowska-Curie ST-IF action (grant agreement No. 101033489 – memory-based percepts: awarded to M. Gandolfo).

## Author Contributions

**Marco Gandolfo:** conceptualization, data curation, data visualization, formal analysis (behavioural and TMS), investigation, methodology, software, visualization, writing – original draft, writing review & editing; funding acquisition.

**Etienne Abassi:** data curation, investigation, formal analysis (fMRI re-analysis), data curation, writing – review and editing, visualization.

**Eva Balgova:** investigation (TMS study), data curation, formal analysis (behavioural and TMS), writing – review and editing, visualisation

**Paul E. Downing:** conceptualization, methodology, software, writing – original draft, writing – review & editing.

**Liuba Papeo:** conceptualization, methodology, resources, writing – review & editing, supervision, funding acquisition.

**Kami Koldewyn:** conceptualization, methodology, writing – original draft, writing – review & editing, supervision, funding acquisition.

## Notes

https://osf.io/m9vp2/

### Competing Interest Statement

The authors have declared no competing interest.

### Summary of Updates

Update after minor revisions

