## Supplemental Materials for "Converging evidence that left extrastriate body area supports visual sensitivity to social interactions"

### fMRI analyses – Extended results with the FBA

Using data from the functional localizer task, for each participant we identified the FBA, separately for each hemisphere by taking the 200 voxels with the highest selectivity for body-stimuli (bodies > faces+objects+places) within an anatomical mask of the fusiform gyrus. A repeated-measures ANOVA with three factors, ROI (EBA/FBA/OPA), hemisphere (Left/Right), and facing direction (facing / non-facing), showed a significant three-way interaction ($F(2,182) = 3.80$; $p = 0.024$; $\eta p2 = 0.04$). Within the FBA-ROI, we found a main effect of hemisphere ($F(1,91) = 12.40$; $p = 0.001$; $\eta p2 = 0.12$), reflecting a stronger effect for body dyads in the right as compared to the left hemisphere, and a main effect of direction ($F(1,91) = 10.52$; $p = 0.002$; $\eta p2 = 0.10$), reflecting a stronger effect for facing as compared to non-facing dyads. We also found a significant hemisphere by facing direction interaction ($F(1,91) = 12.28$; $p < 0.001$; $\eta p2 = 0.12$). There was a larger difference in neural response between facing and non-facing dyads in the left FBA ($t(91) = 4.08$, $p < 0.001$ , $d = 0.43$, Mean difference = 0.33, 95% CI [0.17, 0.49], BF10 = 196), than in the right FBA ($t(91) = 2.16$, $p = 0.034$ , $d = 0.22$, Mean difference = 0.18, 95% CI [0.01, 0.34], BF10 = 1.05).

**Barplots of the behavioural and TMS experiment plotting all the conditions**

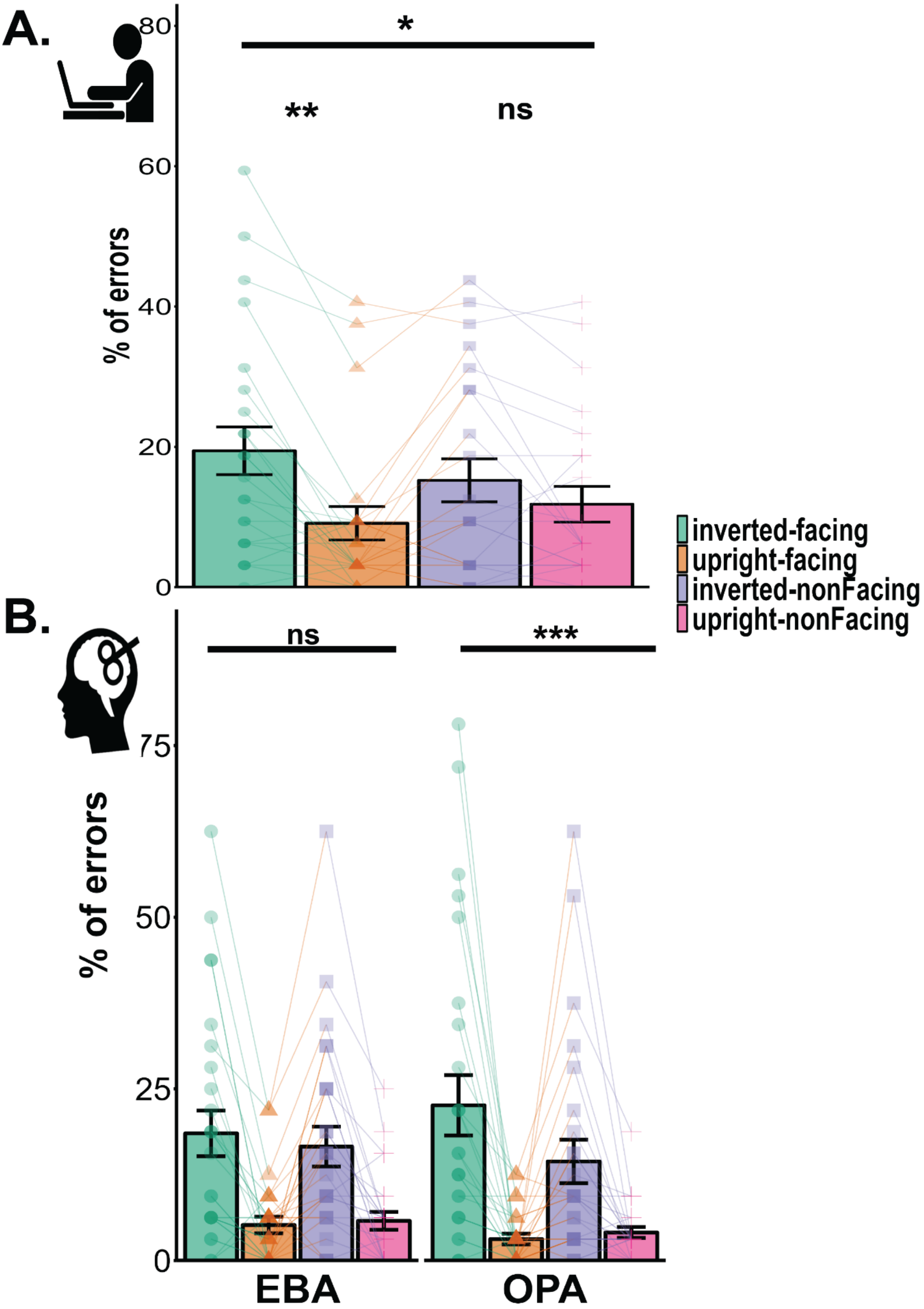


**Figure S1** *Bar chart showing the % of errors for human dyads in each condition. Error bars indicate the standard error of the mean. * p < .05; ** p < .01; *** p < .005.*

**Figures for the chair stimuli:**

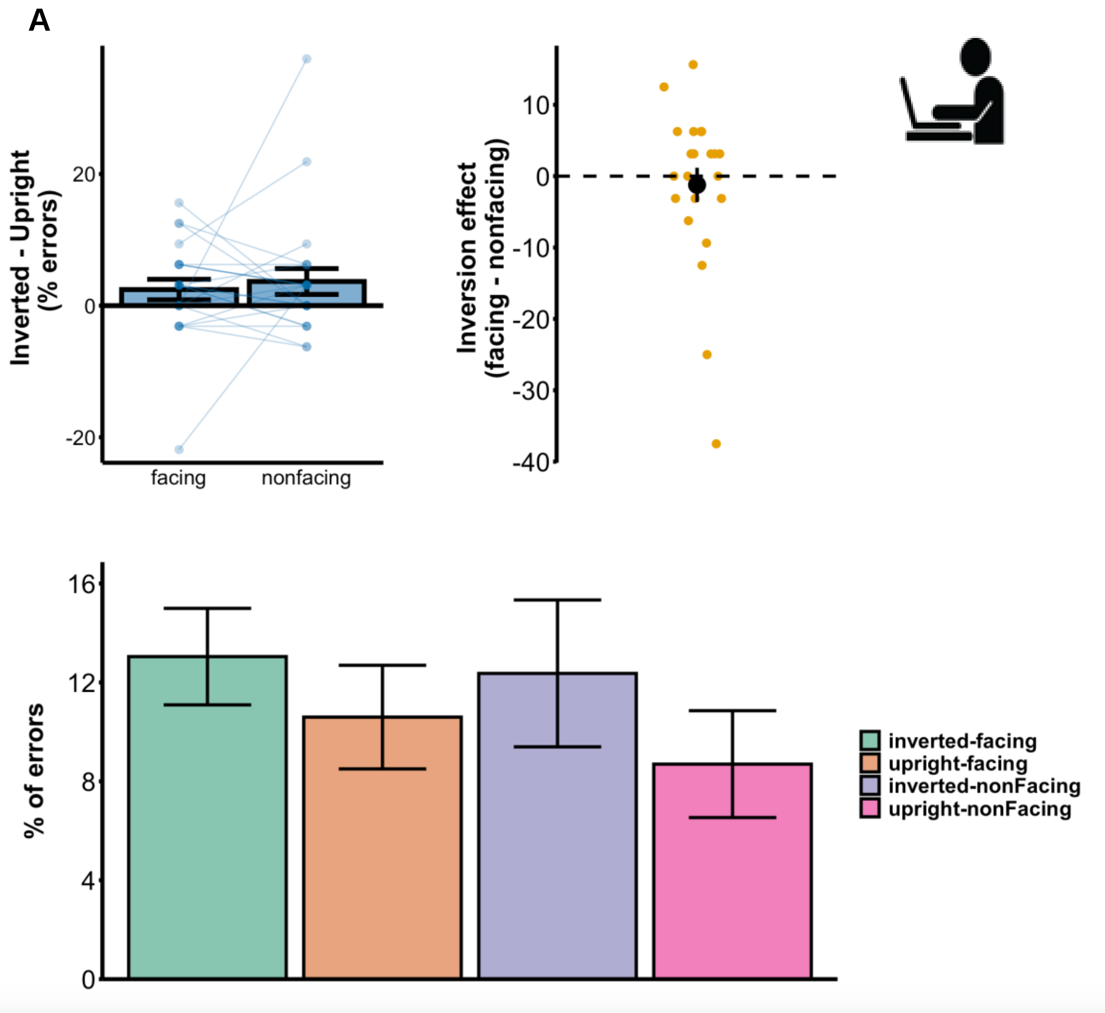


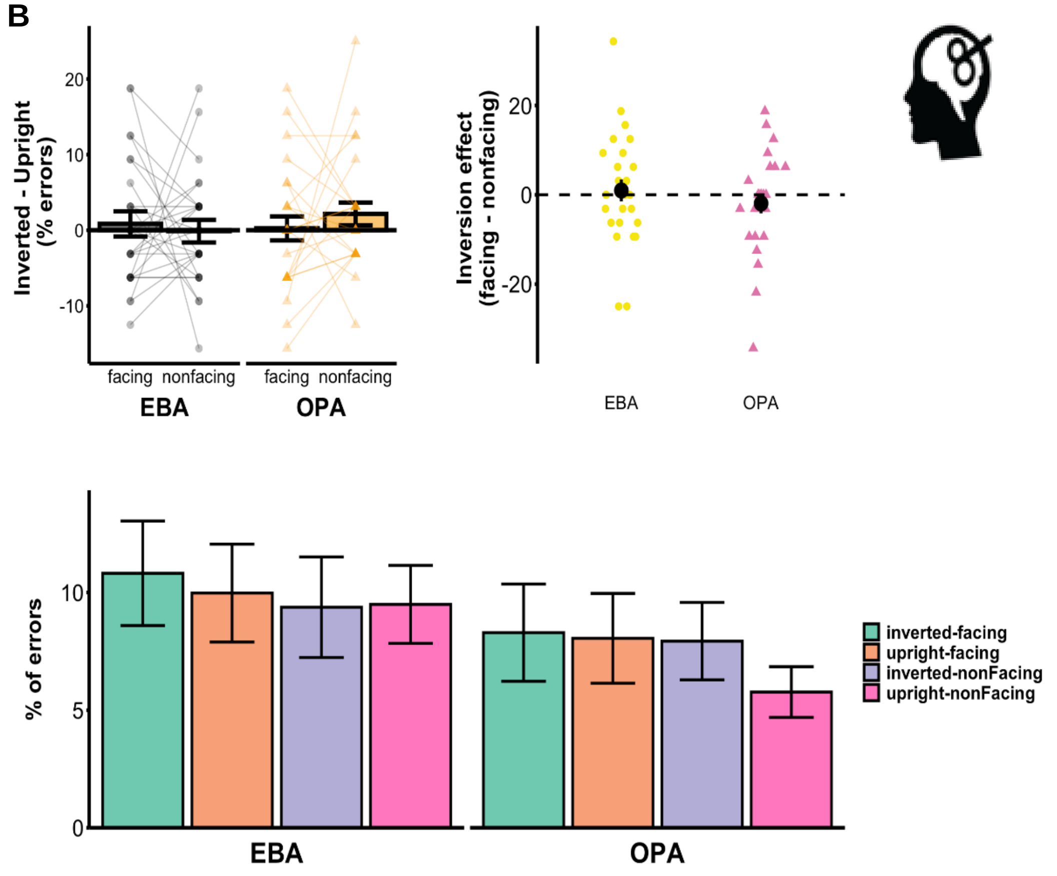

**Figure S2. A.** Results for the chair stimuli on percentage of errors for the behavioural experiment. Top left shows the bar chart of the inversion effect (subtracting the % of errors in the inverted – upright trials). On the right the pointrange plot shows the subtraction of inversion effects with chair stimuli in the facing – non-facing conditions, equivalent index that leads to the 2BIE. At the bottom, the bar for each of the individual conditions of the chair stimuli. **B.** Results for the chair stimuli on percentage of errors for the TMS experiment. Top left shows the bar chart of the inversion effect (subtracting the % of errors in the inverted – upright trials). On the right the pointrange plot shows the subtraction of inversion effects with chair stimuli in the facing – non-facing conditions, equivalent index that leads to the 2BIE. At the bottom, the bar for each of the individual conditions of the chair stimuli in each site. Error bars indicate the standard error of the mean. Points and lines indicate individual data-points for each participant.

**Behavioural study – 2BIE replication – Extended Results**

We observed a main effect of orientation F(1,22) = 21.04, $p < 0.001$, $\eta_p^2 = 0.49$, showing that inversion effects occurred across stimuli and facing direction (upright, M = 15.0, SE = 2.91; inverted, M = 10.1, SE = 2.28). Most importantly, we observed the 3-way interaction indicative of the 2BIE, Stimulus x Orientation x Facing-direction (F(1,22) = 7.43, $p = 0.012$, $\eta_p^2 = 0.25$). The inversion effect for face to face human dyads was larger than for the non-facing dyads (t(22) = 3.014, $p = 0.006$, d = 0.63, Mean Difference = 6.93%, 95% CI [2.16, 11.70], BF10 = 7.16). This difference was absent when contrasting the inversion effect for face-to-face and back-to-back chairs, in which the inversion effect was comparable (t(22) = -0.51, $p = 0.616$, d = -0.11, Mean Difference = 1.22%, 95% CI [-6.21, 3.76], BF10 = 0.25). When contrasting the performance of facing vs nonfacing in each orientation and stimulus, we observed a difference in the inverted human human dyads (t(22) = -2.65, $p = 0.02$, d = 0.55, Mean Difference = 4.21%, 95% CI [0.91, 7.51], BF10 = 3.55,). Performance with the human facing dyad inverted was lower than their non-facing counterpart. No other contrast reached significance (all *p*s > 0.07).

**Reaction times (RTs) and Inverse efficiency scores (ies) analyses**

<u>Behavioural two body inversion effect</u>

*RTs:* The ANOVA on accurate reaction times showed a main effect of orientation (F (1,22) = 20.56, $p < 0.001$, $\eta_p^2 = 0.43$). Participants responded faster to upright (M = 0.50 seconds, SE = 0.02) than inverted trials (M = 0.49 seconds, SE = 0.02) across conditions. We also observed a main effect of category (F(1,22) = 4.604, $p = 0.043$, $\eta_p^2 = 0.17$), in that responses to human dyads were faster (M = 0.49 seconds, SE = 0.02) than responses to chairs (M = 0.50, SE = 0.02). No other effect reached significance (all *p*s > 0.06).

*IES:* The ANOVA on inverse efficiency scores (RT/Proportion of correct responses) showed a main effect of orientation (F (1,22) = 29.75, $p < 0.001$, $\eta_p^2 = 0.58$) and a significant category x orientation interaction (F(1,22) = 6.59, $p = 0.018$, $\eta_p^2 = 0.23$). These main effects were qualified by a significant three-way interaction between category, orientation, and direction (F(1,22) = 6.49, $p = 0.018$, $\eta_p^2 = 0.23$). The inversion effect for face to face human dyads was larger than non-facing human dyads (t(22) = 3.25, *p* = 0.004, d = 0.68, Mean difference = 0.07 seconds, 95% CI [0.02, 0.11], BF10 = 11.5). The inversion effect did not reliably differ instead for the chairs (t(22) = -0.63, *p* = 0.54, d = -0.13, mean difference = 0.01 seconds, 95% CI [-0.06, 0.03], BF10 = 0.26). Therefore, the 2BIE was also present in the inverse efficiency scores.

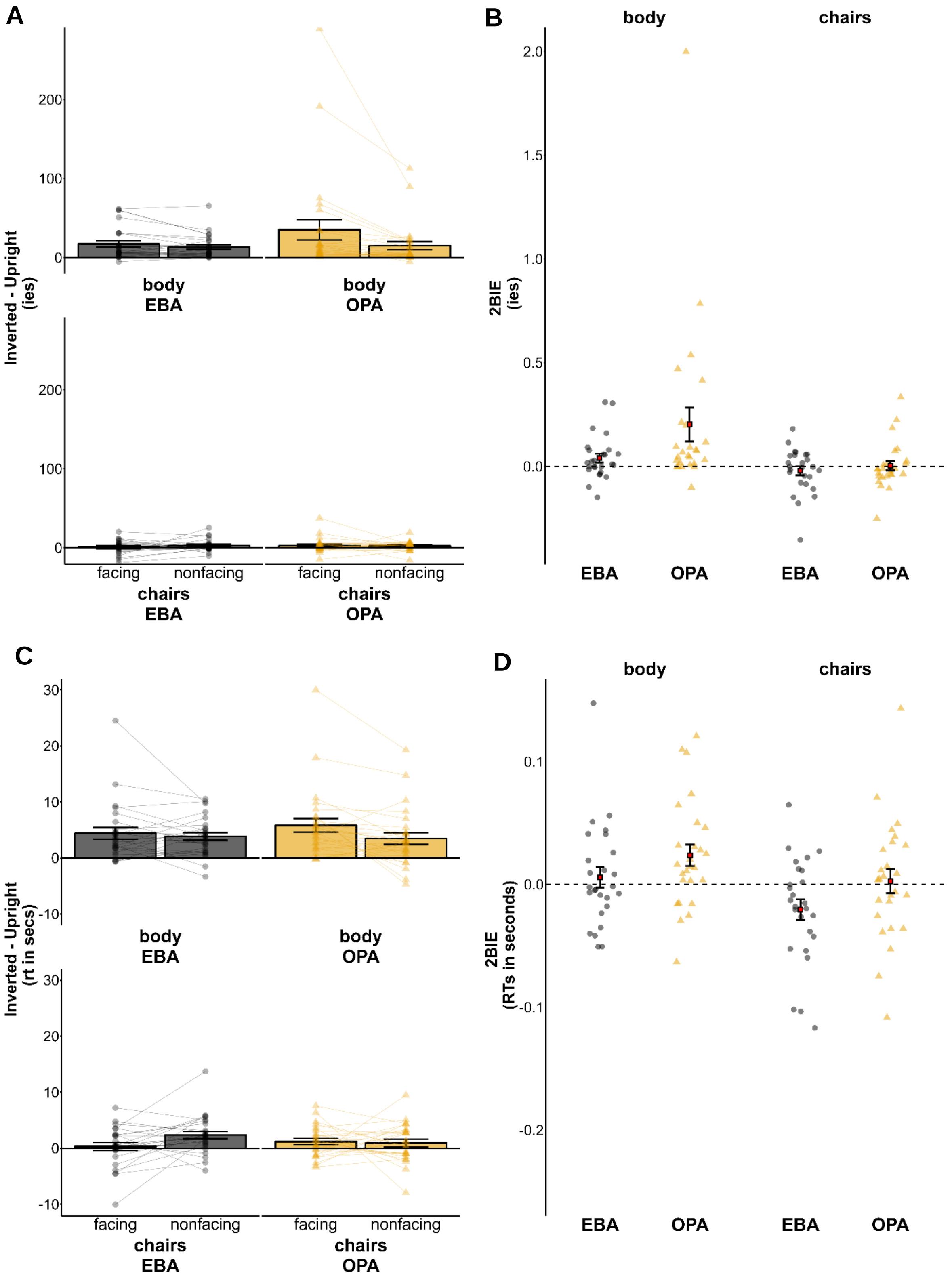

**Figure S3.** The figure shows **A.** the inversion effect (inverted – upright) on the inverse efficiency scores for the chairs and the human dyads after EBA and OPA stimulation. **B.** The difference in inversion effect (the 2BIE) on IES for each stimulus category after EBA and OPA stimulation.
**C.** the inversion effect (inverted – upright) on the accurate reaction times for the chairs and the human dyads after EBA and OPA stimulation. **D.** The difference in inversion effect (the 2BIE) on reaction times for each stimulus category after EBA and OPA stimulation.

<u>TMS</u>

*IES:* Results are shown in Figure **S3A, B**. The ANOVA on inverse efficiency scores showed a main effect of orientation (F (1, 25) = 13.14, $p$ < 0.001, $\eta_p^2$ = 0.34), and a main effect of direction (F (1, 25) = 9.62, $p$ = 0.005, $\eta_p^2$ = 0.28). We also observed a category x orientation interaction (F (1, 25) = 10.83, $p$ = 0.003, $\eta_p^2$ = 0.30), further qualified by a category x orientation x direction interaction (F (1, 25) = 8.18, $p$ = 0.008, $\eta_p^2$ = 0.25). Across stimulation site, the inversion effect reliably differed between facing and non-facing human dyads (t(25) = 2.72, $p$ = 0.012, d = 0.53, Mean Difference = 0.12, 95% CI [0.03, 0.21], BF10 = 4.16), being larger in the facing dyads (M = 0.26, SE = 0.08) than in the non-facing dyads (M = 0.14, SE = 0.04). Conversely, the inversion effect did not differ between facing and non-facing chairs (t(25) = -0.84, $p$ = 0.409, d = 0.16, Mean Difference = 0.01, 95% CI [-0.03, 0.01], BF10 = 0.29). We also observed a site x orientation x direction interaction (F = 4.31, $p$ = 0.048, $\eta_p^2$ = 0.15). Across category, the inversion effect differed between facing and non-facing condition during OPA stimulation (t(25) = 2.38, $p$ = 0.025, d = 0.47, Mean Difference = 0.10, 95% CI [0.01, 0.19], BF10 = 2.19), but not during EBA stimulation (t(25) = -0.64, $p$ = 0.528, d = -0.13, Mean Difference = 0.01, 95% CI [-0.04, 0.02], BF10 = 0.25). The site x category x orientation x direction interaction did not reach significance (F (1, 25) = 2.57, $p$ = 0.12, $\eta_p^2$ = 0.09). Overall, the results on the inverse efficiency scores showed a similar pattern to the percentage of errors, albeit not significantly, ruling out the possibility that our TMS effects on percentage of errors are better explained by speed accuracy trade-offs (see **Figure S3B**).

*RTs:* Results are shown in Figure **S3CD**. The ANOVA on accurate reaction times showed a main effect of category (F (1, 25) = 10.12, $p$ = 0.004, $\eta_p^2$ = 0.29), a main effect of orientation (F (1, 25) = 31.37, $p$ < 0.001, $\eta_p^2$ = 0.56), and a main effect of direction (F (1, 25) = 15.6, $p$ < 0.001, $\eta_p^2$ = 0.38). We also observed a site x orientation x direction interaction (F (1, 25) = 6.93, $p$ = 0.014, $\eta_p^2$ = 0.22). With EBA stimulation, across stimulus category, the inversion effect was not reliably modulated by facing direction (t(25) = -1.50, $p$ = 0.145, d = -0.30, Mean Difference = 0.01 seconds, 95% CI [-0.02, 0.00], BF10 = 0.56). With OPA stimulation, across stimulus category, the effect was modulated by facing direction (t(25) = 2.38, $p$ = 0.03, Mean difference = 0.01 seconds, 95% CI [0.00, 0.02], BF10 = 2.17). We also observed a category x orientation x direction interaction (F (1, 25) = 4.88, $p$ = 0.037, $\eta_p^2$ = 0.16). The inversion effect, across stimulation site, was modulated by facing direction with the human dyads (t(25) = 2.22, $p$ = 0.036, d = 0.44, $p$ = 0.036, Mean Difference = 0.01 seconds, 95% CI [0.01, 0.03], BF10 = 1.65), but not reliably with the chairs (t(25) = -1.46, $p$ = 0.158, d = -0.29, Mean Difference = -0.01, 95% CI [-0.02, 0], BF10 = 0.53). The site x category x orientation x direction interaction did not reach significance (F (1, 25) = 0.08, $p$ = 0.78, $\eta_p^2$ = 0.01).

**Combined analysis for left and right hemisphere stimulation analysis (8 participants)**

This analysis was done to assess, after the COVID-19 lockdown, whether there would have been more chance to find effects (if present) with left- vs right- hemisphere stimulation. Importantly, the overall performance did not reliably differ between the experiments – t(7) = 1.33, $p$ = 0.225, meaning that participants did not overall improve after having completed the first – right stimulation session. We then performed an ANOVA on accuracy scores with those 8 participants that performed both experiments with left and right hemisphere stimulation. Here, we only considered the accuracy scores (proportion of correct responses) on the human dyads and ran a Site x Hemisphere x Orientation x Direction repeated Measure ANOVA. We found a significant interaction among these factors – F(1,7) = 6.14, $p$ = 0.042, $\eta_p^2$ = 0.47). **Figure S6** shows the 2BIE for each hemisphere in each stimulation site. When contrasting the 2BIE against 0, after left EBA stimulation this was not reliable (t(7) = -1.67, $p$ = 0.14, BF10 = 0.91) and numerically was negative (M = -0.02, SD = 0.03). Conversely, after right EBA stimulation this effect was reliable (t(7) = 2.50, $p$ = 0.041, BF10 = 2.20), and numerically positive (M = 0.08, SD = 0.09). The 2BIE was also reliable, and positive, after left OPA stimulation (t(7) = 3.05, $p$ = 0.018, BF10 = 4.05, M = 0.07, SD = 0.07), and numerically positive after right OPA stimulation (t(7) = 1.94, $p$ = 0.093, BF10 = 1.21 – M = 0.07, SD = 0.10). While conclusions on these data should be taken carefully due to the low power, they gave a clear hint that finding a reduced 2BIE after left EBA stimulation would have been more likely than after the right hemisphere stimulation. This was in line with the results we obtained from the fMRI re-analysis that was ran in parallel. This analysis showed indeed a left hemispheric preference for facing vs non-facing human dyads.

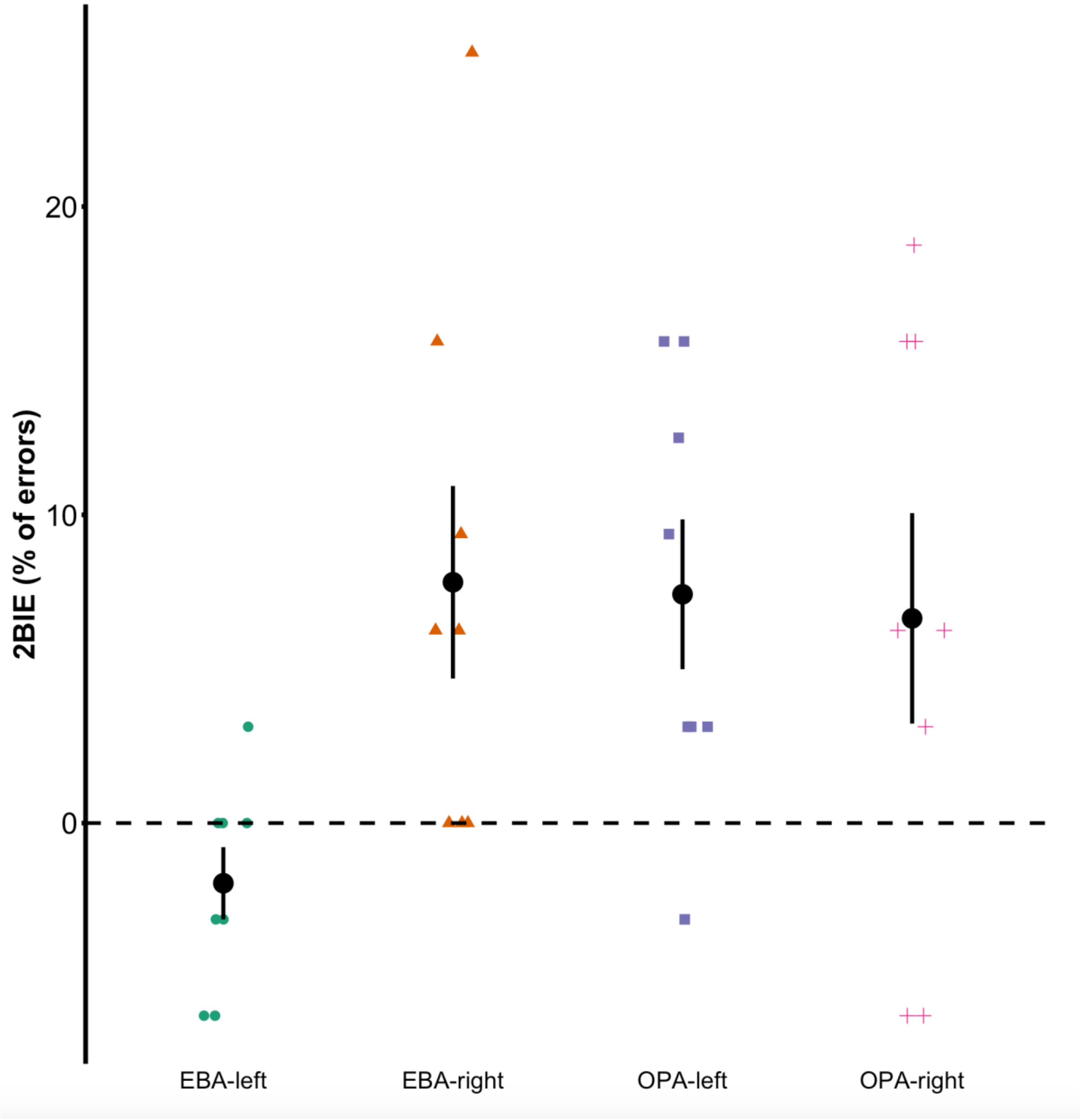


**Figure S4.** The point-range plot shows the 2BIE in each hemisphere and stimulation site for the 8 participants who completed both experiments.

**Right Hemisphere stimulation analysis on (12 Participants)**

The ANOVA on the accuracy scores (proportion of correct responses) did not show the highest-level interaction i.e. Site x category x orientation x facing-direction ($F(1,11) = 0.002$, $p = 0.97$, see **Figure S5**).

An ANOVA on the accuracy scores showed a main effect of orientation ($F(1,11) = 18.52$, $p < 0.001$, $\eta_p^2 = 0.14$). Performance was higher in the upright (M = 0.95, SD = 0.07) than in the inverted trials (M = 0.87; SD = 0.13). Further, there was a category x orientation x facing direction interaction ($F = 8.99$, $p = 0.01$, $\eta_p^2 = 0.45$). This interaction replicates the pattern found in the behavioural experiment – i.e. that the inversion effect was larger with human facing dyads than in the other conditions (**Figure S7**). However, after splitting by category, the ANOVA on human dyads did not show a significant orientation x direction interaction ($F(1,11) = 4.28$, $p = 0.063$, $\eta_p^2 = 0.28$), likely due to lack of power to detect such effect. No other effect reached significance (all *p*s > 0.076).

**Table S1.** Individual MNI coordinates for functionally localised left and right EBA.

| subject | site | x | y | z |
|---|---|---|---|---|
| 1 | EBA | -47.1 | -71.8 | 6.2 |
| 1 | OPA | -33.9 | -81.6 | 27 |
| 2 | EBA | -49.5 | -74.9 | 8.3 |
| 2 | OPA | -30.5 | -88.9 | 20.2 |
| 3 | EBA | -43.8 | -75 | -1.3 |
| 3 | OPA | -23 | -87.9 | 17.6 |
| 4 | EBA | -46.7 | -68 | 3.7 |
| 4 | OPA | -28.7 | -80.9 | 17.6 |
| 5 | EBA | -38 | -81.6 | 7.1 |
| 5 | OPA | -28.3 | -80.9 | 28.4 |
| 6 | EBA | -46.3 | -70.4 | -2.4 |
| 6 | OPA | -28.9 | -80.7 | 3.2 |
| 7 | EBA | -46.1 | -76.4 | -7.3 |
| 7 | OPA | -34 | -85.1 | 21.9 |
| 8 | EBA | -47 | -63.8 | 10.2 |
| 8 | OPA | -28.3 | -82.3 | 29.4 |
| 9 | EBA | -38.5 | -82.3 | 2.6 |
| 9 | OPA | -35.4 | -78.4 | 19.2 |
| 10 | EBA | -51 | -74.4 | 6.6 |
| 10 | OPA | -32.4 | -81.4 | 21.6 |
| 11 | EBA | -50.3 | -75.3 | -1.1 |
| 11 | OPA | -33.8 | -83.5 | 16.9 |
| 12 | EBA | -40.7 | -73.3 | 2 |
| 12 | OPA | -29 | -80 | 23.1 |
| 13 | EBA | -44.3 | -67.8 | -7 |
| 13 | OPA | -28.3 | -84.9 | 22.1 |
| 14 | EBA | -34.8 | -87.2 | 4.7 |
| 14 | OPA | -33.1 | -88.9 | 20.1 |
| 15 | EBA | -40.3 | -65.1 | -8.1 |
| 15 | OPA | -29.5 | -84.9 | 13 |
| 16 | EBA | -42.3 | -85.9 | -15.2 |
| 16 | OPA | -35.9 | -83.1 | 5.5 |

| | | | | |
|---|---|---|---|---|
| 17 | EBA | -46.9 | -61.7 | -9.9 |
| 17 | OPA | -23.2 | -85.6 | 17.9 |
| 18 | EBA | -39.6 | -73.7 | -13.4 |
| 18 | OPA | -32 | -86.1 | 11.4 |
| 19 | EBA | -49.5 | -71.5 | 13.8 |
| 19 | OPA | -34.8 | -80.1 | 6.2 |
| 20 | EBA | -52.5 | -62.1 | -3.9 |
| 20 | OPA | -37.1 | -82.3 | 10.9 |
| 21 | EBA | -37.8 | -70.9 | -16.4 |
| 21 | OPA | -32.9 | -84.3 | 10.1 |
| 22 | EBA | -44.6 | -73.1 | -11.9 |
| 22 | OPA | -39.1 | -88.6 | 1.3 |
| 23 | EBA | -52.8 | -66.8 | -8.4 |
| 23 | OPA | -35.9 | -81.2 | 20.9 |
| 24 | EBA | -48.7 | -65.2 | 12.3 |
| 24 | OPA | -31.9 | -80.8 | 23.6 |
| 25 | EBA | -37.7 | -77.1 | -2.7 |
| 25 | OPA | -29.5 | -85.5 | 11.1 |
| 26 | EBA | -43.1 | -75.6 | -9.3 |
| 26 | OPA | -29.6 | -88.8 | 9.6 |
